# Coping with Pleistocene climatic fluctuations: demographic responses in remote endemic reef fishes

**DOI:** 10.1101/672774

**Authors:** Erwan Delrieu-Trottin, Nicolas Hubert, Emily C. Giles, Pascaline Chifflet-Belle, Arnaud Suwalski, Valentina Neglia, Cristian Rapu-Edmunds, Stefano Mona, Pablo Saenz-Agudelo

## Abstract

Elucidating demographic history during the settlement of ecological communities is crucial for properly inferring the mechanisms that shape patterns of species diversity and their persistence through time. Here, we used genomic data and coalescent-based approaches to elucidate for the first time the demographic dynamics associated with the settlement by endemic reef fish fauna of one of the most remote peripheral islands of the Pacific Ocean, Rapa Nui (Easter Island). We compared the demographic history of nine endemic species in order to explore their demographic responses to Pleistocene climatic fluctuations. We found that Rapa Nui endemic species share a common demographic history as signatures of population expansions were retrieved for almost all of the species studied here, and synchronous demographic expansions initiated during the last glacial period were recovered for more than half of the studied species. These results suggest that eustatic fluctuations associated with Milankovitch cycles have played a central role in species demographic histories and in the final stage of the community assembly of many Rapa Nui reef fishes. Specifically, sea level low stands resulted in the maximum reef habitat extension for Rapa Nui endemic species; we discuss the potential role of seamounts in allowing endemic species to cope with Pleistocene climatic fluctuations, and we highlight the importance of local historical processes over regional ones. Overall, our results shed light on the mechanisms by which endemism arises and is maintained in peripheral reef fish fauna.

## 1 INTRODUCTION

Demographic expansions are required to colonize novel environments and to establish new populations. Tracking the direction (expansions or contractions) and the magnitude of demographic changes through time can help elucidate not only when populations were established, but also reveal the factors that triggered such changes. For instance, if abiotic factors determined population sizes, we would expect to see populations expand or contract as environment changes. In contrast, population sizes should remain constant or decoupled from environmental changes when adaptation to new conditions is possible (Jump & Penuelas, 2005; Vrba, 1992; Wong & Candolin, 2015). Populations of different species can vary in their abilities to colonize novel environments and in the way that they cope with environmental variability (Hewitt, 2001, 2004; Lessa, Cook, & Patton, 2003). The degree to which colonization dynamics and/or responses to environmental changes are synchronous have not been well-characterized. Overall, uncovering patterns acting at multispecies levels is crucial to understanding how climatic fluctuations can ultimately drive patterns of community assembly and thus the distribution of current biodiversity (Chan, Schanzenbach, & Hickerson, 2014).

Colonization dynamics and responses to climatic fluctuations can leave a detectable genetic footprint of changes in population size (Wakeley, 2009, 2010). A large number of studies have shown that past major climatic fluctuations, with expansions and contractions of ice sheets and sea level fluctuations (Cutler et al., 2003; Pahnke, Zahn, Elderfield, & Schulz, 2003) have indeed deeply impacted the demography of many terrestrial and marine species (Avise, 2009; Hewitt, 2003; Hickerson et al., 2010; Petit et al., 2003). Theory predicts that assemblages of co-distributed species with similar physiological and habitat requirements should respond similarly to environmental changes (Hewitt, 2001, 2004b; Lessa, Cook, & Patton, 2003). In the same manner, assemblages of co-distributed species of similar ages should display similar colonization timings and dynamics, as well as concerted changes in their population sizes through time. Synchronicity of population size changes of co-distributed taxa has only been recovered a few times in terrestrial and marine environments (e.g. Burbrink et al., 2016; Chan et al., 2014; Reid, Naro-Maciel, Hahn, FitzSimmons, & Gehara, 2019).

Coral reefs are among the most diverse ecosystems, yet how this biodiversity has arisen and is maintained remains largely debated. Among coral reef organisms, coral reef fishes represent the most diverse and widely distributed group of vertebrates on the planet (Nelson, Grande, & Wilson, 2016). Their geographic distribution is uneven; in particular, endemism hotspots, i.e. regions hosting high proportions of species with geographically restricted distributions, are found along the periphery of the Indo-Australian Archipelago (Allen, 2008; Bellwood & Wainwright, 2002; Delrieu-Trottin et al., 2015; Eschmeyer, Fricke, Fong, & Polack, 2010; Gaboriau, Leprieur, Mouillot, & Hubert, 2018; Hughes, Bellwood, & Connolly, 2002; Randall, 2005, 2007; Randall & Cea, 2011). As such, a series of questions remain unanswered regarding the mechanisms enabling the co-existence of hundreds of species locally and the existence of large diversity gradients in the Indo-Pacific (Allen & Erdmann, 2012). Reconstructing the demographic histories of populations in these hotspots of endemism could provide key information on how coral reef fish biodiversity is maintained in coral reef ecosystems.

Rapa Nui (Chile, 166 km2) is the most remote peripheral hotspot of endemism of reef fishes; it is composed of 21.7% endemic species and includes all major reef fish families and a high diversity of life history traits (Cea, 2016; Randall & Cea, 2011). Despite being the second, after Hawai’i (Randall, 1998, 2007), hotspot of endemism for reef fishes in the tropical Indo-Pacific, the mechanisms that have underlined the persistence of species with such range-restricted distributions through geological time remain enigmatic. The Rapa Nui reef fish community is not only exceptional for its high percentage of endemic species, but also for being extremely species-poor for a tropical reef system (only 169 fish species; 139 shore fishes) By comparison, the Indo-Australian Archipelago houses more than 2,600 reef fish species (Allen & Erdmann, 2012). Rapa Nui and the islet Motu Motiro Hiva (Salas y Gómez, 0.15 km2), located 400 km further east and which shares the same reef fish fauna as Rapa Nui (Friedlander et al., 2013), constitute the only two emerged islands of the Easter Chain. Rapa Nui and Motu Motiro Hiva are relatively young; 2.5 and 1.7 My, respectively (Clouard & Bonneville, 2005), but they are embedded in a network of numerous seamounts. This seamount chain extends 2,232 km east to the Nazca seamount (23°360 S, 83°300 W) (Clouard & Bonneville, 2005; Ray et al., 2012) and these mounts have emerged to various degree during periods of low sea level. In this context, an “Ancient Archipelago” hypothesis has been formulated by Newman & Foster (1983) to account for the high level of endemism observed in Rapa Nui and Motu Motiro Hiva that further questions the dynamics of community assembly in such a remote system. This hypothesis states that seamounts could have provided potentially suitable habitat for the past 29 My during periods of low sea level (lowstands) for endemics of the region, allowing them to evolve and persist up to present times (Newman & Foster, 1983). Interestingly, a recent analysis of the divergence times of endemic species from their closest relatives indicates that most species endemic to Rapa Nui are neoendemics, i.e. younger than the emergence of Rapa Nui and Motu Motiro Hiva (Delrieu-Trottin et al., 2019).

Seamounts near Rapa Nui could have played a major role in species persistence through time, especially during the sea-level fluctuations of the Pleistocene that resulted in changes of up to 150 m below present levels (Miller et al., 2005). This hypothesis is increasingly called to attention as recent surveys showed that many Rapa Nui endemic species can be found from shallow waters to the mesophotic zone (at least 160 m) in Rapa Nui waters and in nearby seamounts (Easton et al., 2017). The paleo-environmental perturbations of the Pleistocene might be expected to have left a footprint on the community assembly and the associated demographic histories of these populations (Bard et al., 1996; Delrieu-Trottin et al., 2017; Hewitt, 2004; Ludt & Rocha, 2015; Woodroffe et al., 2010). For instance, numerous species are reported as present in Rapa Nui waters but are actually vagrants, i.e. species that colonize Rapa Nui from time to time without being able to establish a stable population locally (Randall & Cea, 2011), suggesting the current isolated state of Rapa Nui is not well-suited for species immigration compared to lowstands. Reconstructing the demographic histories of populations in this hotspot of endemism could provide key information on how endemic species cope with Pleistocene climatic fluctuations.

The present study investigates the demographic history of multiple species of coral reef fishes in Rapa Nui possessing distinct range sizes and life-history traits. We used genome-wide sequencing (ddRAD) and coalescent-based approaches in order to highlight potential demographic trends during community assembly. We hypothesize that sea-level fluctuations have left different signatures on the demographic history of Rapa Nui reef fish species, and these depend on specific characteristics of species. In particular, (1) we expect that species with large range distributions have functioned as a metapopulation and that this metapopulation dynamics have mitigated the impacts of sea level fluctuations on their demographic histories, (2) fish species restricted to shallow waters should have more frequent fluctuations of population sizes; thus, wider occurrence along a depth gradient could mitigate the impact of sea level fluctuations. Lastly (3) signals of population growth in Rapa Nui endemic fishes should have occurred during low stands. This is expected as a corollary of the Ancient Archipelago hypothesis where the higher number of emerged seamounts around Rapa Nui should have favored population expansions. To our knowledge, this represents the first attempt to reconstruct the demographic history of an assemblage of endemic coral reef fishes of one of the least studied coral reef communities of the Indo-Pacific Ocean.

## 2 MATERIALS AND METHODS

### 2.1 Sampling

Rapa Nui hosts two types of endemic species, small-range and large-range endemic species. Small-range endemics are only present around Rapa Nui and Motu Motiro Hiva and have a maximum range of <500 km in linear distance, see Delrieu-Trottin, Maynard, & Planes (2014). Southern subtropical endemics have large-ranges (1,000– 8,000 km in linear distance, see Delrieu-Trottin et al. (2014), hereafter large-range endemic species) and are distributed from Southern Polynesia to Rapa Nui (regional endemics in Friedlander et al. (2013)). Large-range endemics, contrary to small-range endemics, are present both in Rapa Nui and in other southern subtropical islands of the Pacific. As such, populations of these fishes are embedded in metapopulations of larger geographic ranges than those of small-range endemics, making them potentially prone to not only processes at the scale of Rapa Nui (local processes) but also at the scale of the South Pacific (regional processes). A total of 143 reef fishes, belonging to nine species, were collected using polespears or an anesthetic (clove oil) in Rapa Nui in October 2016 (**TABLE 1**). The fishes collected represent the two types of endemics (small-range endemics: 7 species; large-range endemics: 2 species). The species analyzed here belong to seven major reef fish families and possess different reproductive strategies, with five species producing pelagic eggs, three species producing demersal eggs, and one species brooding eggs in their mouths (**TABLE 1**). Overall, we investigated the consequences of historical processes, such as sea level changes, and ecological dynamics, i.e. those driven by reproductive strategy, on the observed genetic diversity. If historical processes were more important than ecological processes in the maintenance of endemic species in Rapa Nui, we would expect to find similar population dynamics for the two types of endemic species.

**TABLE 1.** Ecological data on the species of interest of this study. Range size (S: small-range size; L: large-range size), family, reproductive strategy (p: pelagic eggs; b: demersal eggs; m: mouth brooding), maximum depth where the species have been observed (Max. depth) (1 to 5); generation time used for this study (6 to 9); maximum size of the species (Max. size) and references associated (^1^: specimens observed freediving by Cristian Rapu-Edmunds; ^2^: Randall & Cea, 2011; ^3^: specimens video-recorded at those depths (pers. comm. Erin Easton); ^4^: Easton et al., 2017 ^5^: specimens photographed at those depths by com. pers Luiz A. Rocha in 2016; ^6^: Craig, Eble, Bowen, & Robertson, 2007, ^7^: Bowen, Muss, Rocha, & Grant, 2006, ^8^: Craig et al., 2010, ^9^: Gajdzik, Bernardi, Lepoint, & Frédérich, 2018).

| Range size | Family | Reproductive strategy | Species | Max. depth (m) | Generation time (years) | Max. size (TL, cm) |
| --- | --- | --- | --- | --- | --- | --- |
| L | Holocentridae | pelagic eggs | *Myripristis tiki* | 80<sup>1</sup> | 3<sup>6</sup> | 26.5<sup>2</sup> |
| L | Pomacanthidae | pelagic eggs | *Centropyge hotumatua* | 50<sup>2</sup> | 1<sup>7</sup> | 9<sup>2</sup> |
| S | Apogonidae | mouthbrooding | *Ostorhinchus chalcus* | 25<sup>2</sup> | 2 | 16<sup>2</sup> |
| S | Chaetodontidae | pelagic eggs | *Chaetodon litus* | 156<sup>3</sup> | 2<sup>8</sup> | 15.5<sup>2</sup> |
| S | Holocentridae | pelagic eggs | *Sargocentron wilhmi* | 157<sup>4</sup> | 3<sup>6</sup> | 19.5<sup>2</sup> |
| S | Labridae | pelagic eggs | *Coris debueni* | 70<sup>2</sup> | 3 | 27<sup>2</sup> |
| S | Monacanthidae | benthic eggs | *Cantherhines rapanui* | 20<sup>2</sup> | 3 | 20<sup>2</sup> |
| S | Pomacentridae | benthic eggs | *Chrysiptera rapanui* | 75<sup>2,3</sup> | 1<sup>9</sup> | 7.8<sup>2</sup> |
| S | Pomacentridae | benthic eggs | *Chromis randalli* | 105<sup>5</sup> | 2<sup>9</sup> | 15<sup>2</sup> |

### 2.2 Library preparation and sequencing

Whole genomic DNA was extracted from fin or gill tissue preserved in 96% ethanol using the GeneJet Genomic DNA purification kit according to the manufacturer’s protocols (Thermo Fisher Scientific). Double-digest restriction-associated DNA (ddRAD) libraries were prepared following Peterson et al.’ (2012) using EcoRI and MspI restriction enzymes, a 400 bp size selection and a combination of two indexes and 24 barcodes to pool 48 individuals per lane. The genomic libraries obtained were sequenced in three lanes of a HiSeq 2500 Illumina sequencer (single end, 125 bp). Illumina reads are available from the Sequence Read Archive (SRA) at NCBI under the BioProject Accession no: PRJNA622670.

### 2.3 De novo assembly and SNP calling

We used the ‘process_radtags.pl’ pipeline in STACKS version 2.0 (Catchen, Hohenlohe, Bassham, Amores, & Cresko, 2013; Catchen, Amores, Hohenlohe, Cresko, & Postlethwait, 2011) to demultiplex and quality filter the sequences obtained. In the absence of reference genomes for the species under study, RADSeq loci were assembled de novo using the ‘denovo_map.pl’ pipeline in STACKS each species separately. We used the parameter combination recommended by Mastretta-Yanes et al. (2015); this included minimum read depth to create a stack (m) = 3, number of mismatches allowed between loci within individuals (M) = 3, number of mismatches allowed between loci within catalogue (n) = 3 and required a locus to be present in all individuals of each species (r = 1). Following de novo mapping, an initial data-filtering step was performed using the population component of STACKS removing all loci with maximum observed heterozygosity higher than 0.8. We kept all single-nucleotide polymorphisms (SNP) per stack (i.e. locus) and did not use any threshold regarding the minor allele frequencies as these rare variants are informative when performing demographic inference (Wakeley, 2009). We then removed all loci displaying more than three SNPs to avoid potential paralogues. Heterozygosity (H), FIS, mean pairwise differences (π) computed on all sites of variable loci and Watterson’s theta (*Θ*_*w*_) computed on all sites of variable loci were calculated in R (R Core Team, 2017) using the packages vcfR (Knaus & Grünwald, 2017), adegenet (Jombart, 2008; Jombart & Ahmed, 2011), PopGenome (Pfeifer, Wittelsbürger, Ramos-Onsins, & Lercher, 2014) and pegas (Paradis, 2010).

### 2.4 Statistical analyses of genetic diversity indices and reconstruction of potential habitats given present and past sea level conditions

We tested whether the summary statistics (*Θ*_*w*_, π, He), neutrality tests (Tajima’s D and Fu and Li’s D) and TMRCA obtained for each species were related to range size (small-range / large-range endemics), and/or reproductive strategies (pelagic eggs / demersal eggs / mouth brooding). We used a multivariate regression tree (MRT) (De’ath, 2002) on normalized summary statistics to hierarchize the significant predictor variables. Prediction error was used to assess model fit and determine the appropriate tree size, and the tree was pruned by cross-validation using the minimum rule (Breiman, Friedman, Olshen, & Stone, 1984). All statistical analyses were performed in R, using the packages vegan (Oksanen et al., 2012) and mvpart (Therneau, Oksanen, Oksanen, Atkinson, & De’ath, 2014) while graphical representations were performed using ggplot2 (Wickham, 2009).

Finally, as Rapa Nui and Motu Motiro Hiva are part of the Easter chain composed of numerous seamounts, we examined how eustatic changes modified the surface of potential habitat for Rapa Nui endemic species. We investigated whether periods corresponding to minimum sea level constituted an extension or a reduction of potential available habitat for Rapa Nui endemic fishes using the package marmap (Pante & Simon-Bouhet, 2013) in R. We reconstructed both present surface habitat using the maximum depth recorded for all species (160 m) and surface habitat given the scenario of a minimum sea level (sea level lowstand, -120 m context), corresponding to the surface between -120 and -280 m depth.

### 2.5 Demographic inferences

To detect departures from the neutral Wright–Fisher model, we computed Tajima’s D and Fu & Li’s D neutrality tests implemented in PopGenome. Significance of Tajima’s D (TD) was evaluated after 1000 coalescent simulations of a constant population model with size *Θ*_*w*_ using fastsimcoal and a custom script. Significant negative values of these tests indicate population growth while significant positive values are a signature of either genetic subdivision or population contraction assuming selective neutrality. The folded Site (Allele) Frequency Spectrum (SFS), describing the distribution of allele frequencies across polymorphic sites, was computed in R with the package pegas (Paradis, 2010). The SFS represents a summary of genetic variation whose shape is sensitive to underlying population genetic processes (population size change, migration, selection, etc.). The aggregated SFS (aSFS, Xue & Hickerson, 2015) was then computed using a custom R script. The aSFS combines an array of independent SFSs computed in each species into a single SFS vector. First, the independent SFSs need to have the same number of classes (i.e., each species must have the same number of individuals). To this end, we downsampled all SFSs to 13 diploid individuals, the minimum sample size in our nine species. Each SFS was then transformed into a proportion vector by dividing each class by the total number of SNPs. We then arranged the nine bins within each frequency class in descending order of proportion of total SNPs. The ordering is therefore independent in each frequency class, so that there is no relationship between the initial order of the single taxon and the resulting aSFS, which guarantees exchangeability across single taxon SFSs. The final vector contains a number of classes equal to the number of species times the number of frequency classes (i.e., 9 × 13 in our case).

Demographic inferences were investigated reconstructing the variation in the effective population size (Ne) through time using three independent approaches. First we ran the composite likelihood approach implemented in the software stairwayplot (Liu & Fu, 2015). The *stairwayplot* is a non-parametric model where Ne is free to vary at each coalescent interval. The composite likelihood is evaluated as the difference between the observed SFS and its expectation under a specific demographic history.

Second, we ran an approximate Bayesian computation algorithm based on coalescent simulations following Maisano Delser et al (2016) to compare the results obtained by the *stairwayplot*. Briefly, we performed 1,000,000 coalescent simulations of a demographic model with three instantaneous changes of Ne. This number was chosen as a compromise between over parametrization and the need to capture multiple demographic changes through time. We further ran a simplified model with only one instantaneous size change to determine if its use in the hierarchical ABC analysis (hABC, see below) would have failed to detect important historical events and biased the results. The full model is therefore defined by seven parameters: four values of Ne and three instantaneous time changes (hereafter, T). We set the same prior uniform distribution for the four Ne values and incremental uniform distribution for the three T parameters, similarly to Maisano Delser et al. (2016). The simplified model is defined by three parameters: two values of Ne and one instantaneous size change. The two models provided similar results and we therefore used the simplified one in the hABC. Coalescent simulations and SFS computation were performed with fastsimcoal (Excoffier, Dupanloup, Huerta-Sánchez, Sousa, & Foll, 2013a). We used the SFS and the mean pairwise differences (π) as summary statistics. We retained the best 5,000 simulations to perform a local linear regression (Beaumont, Zhang, & Balding, 2002) and reconstructed the abc skyline at user specified time points as in Maisano Delser et al. (2016).

Third, we performed a hierarchical approximate Bayesian computation (hABC) analysis with the goal of detecting potential concerted demographic histories across the species under study (Chan et al., 2014). This method allows combining the datasets from the nine species investigated here into a single analysis in order to estimate objectively when population size changes occurred, whether they were synchronous, and the timing of such demographic changes (Burbrink et al., 2016; Chan et al., 2014). We first extracted the number of species *ζ* with synchronous changes in effective population size from a uniform prior distribution bounded between 1 and n (total number of species). We then extracted the time of co-change (τ_s_) and attributed this to *ζ* randomly chosen species. If *ζ* < n, the remaining times were independently extracted from the same prior used for τ_s_. We then computed E(τ), the average over the n times of change, and the dispersion index computed as the ratio between the variance of these times and E(τ) (Chan et al., 214). Each species follows a simple model of instantaneous size change with two Ne (modern and ancestral) extracted from independent prior distributions (i.e., two priors for each species). We defined the same priors for both Ne to avoid weighting expansions or contractions *a priori*. The rationale here is that some species could have started expanding while others were contracting at the same time (for example, species adapted to cold climate will contract when species adapt to warmer weather will expand, having synchronous though opposite responses). We also took species specific generation times into account when running the simulations. We first performed hABC on the nine species and then restricted the analysis to the species displaying the most similar demographic profiles (see Results). We performed 2,000,000 coalescent simulations with fastsimcoal (Excoffier et al., 2013a) and computed summary statistics (aSFS, mean and standard deviation over species of *Θ* estimated from singleton, Π, and TD) with a custom R script. Local linear regression was finally applied to the best 5,000 simulations to estimate the posterior distribution of each parameter. We performed a cross validation experiment to determine whether we could correctly infer *ζ* given our settings (number of species, loci and summary statistics) by randomly sampling 1,000 datasets generated from prior distributions and re-analyzed them under the same hABC conditions. Finally, a posterior predictive test was run by sampling 50,000 random combinations of parameter values from the hABC posterior distributions with which we simulated pseudo-observed datasets (Bertorelle et al. 2010). A principal component analysis was then applied to check if the estimated model could reproduce the observed data by comparing the aSFS computed using the pseudo-observed and the observed data.

Uncertainty in both mutation rates and generation times can potentially bias molecular dating. Estimated mutation rates for SNP data of fish range from 2.5 × 10^−8^ (Kavembe, Kautt, Machado-Schiaffino, & Meyer, 2016) to 3.5 × 10^−9^ (Malinsky et al., 2018) and their selection for estimating demographic histories is not often justified. As such, for all analysis here we chose the hypothesized fish SNP mutation rate (µ) most often used in the literature; 1.0 × 10^−8^ / site / generation (Jacobs, Hughes, Robinson, Adams, & Elmer, 2018; Le Moan, Gagnaire, & Bonhomme, 2016; Rougeux, Bernatchez, & Gagnaire, 2017; Souissi, Bonhomme, Manchado, Bahri-Sfar, & Gagnaire, 2018; Tine et al., 2014). Estimations of generation times in the wild for fishes are scarce. Most of the generation times we used (six out of nine) were selected from the literature from species that are phylogenetically closely-related to the species studied here (same genus) (**TABLE 1**). Alternatively, we inferred generation time using maximum standard length (three species), as maximum standard length and generation time are positively correlated (Froese & Binohlan, 2000).

## 3 RESULTS

### 3.1 Raw sequence filtering, assembly, and SNP calling

A total of 499,403,262 reads were obtained for the 143 individual samples from the nine species endemic to Rapa Nui. The different filtering steps resulted in the building of an average of 41,470 RAD loci of 120 bp per species (min: 25,331 loci (*Chrysiptera rapanui*); max: 63,590 loci (*Ostorhinchus chalcius*)) out of which an average of 23,693 loci were variable. *Myripristis tiki* displayed the highest number of variable RAD seq loci (28,855) while *Chrysiptera rapanui* had the lowest (19,246) (**TABLE 2**). *Chrysiptera rapanui* displayed the highest percentage of variable RAD seq loci (76%) while *Ostorhinchus chalcius* displays the lowest (38%). The variable RAD loci harbored an average of 37,107 SNPs (min: 28,239 (*Cantherhines rapanui*); max: 46,718 (*Myripristis tiki*)). The depth of coverage per SNP ranged from 31x to 50x (43x averaged across species).

**TABLE 2.**
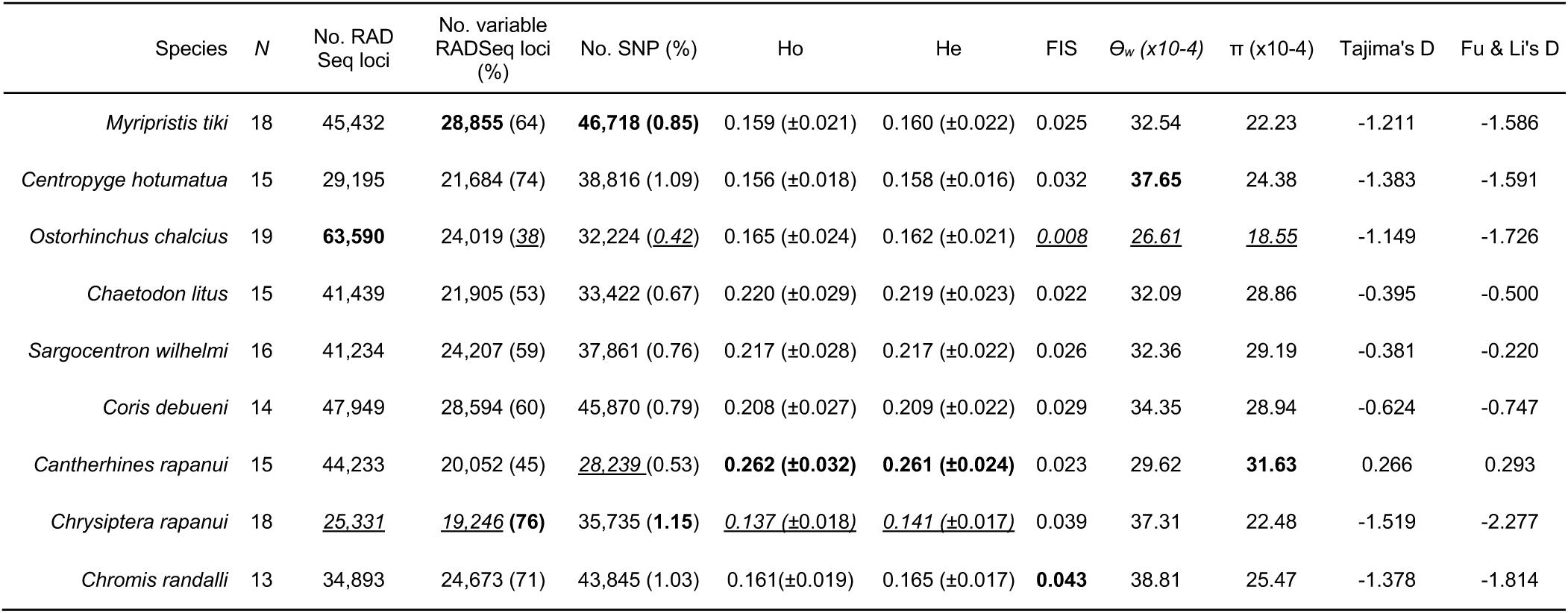
Summary of genetic data for each species. Sample size (*N*) and molecular metrics for each species of the study. Observed (Ho) and expected (He) heterozygosity computed on all sites of variable loci; Watterson’s theta (*Θ*_*w*_) computed on all sites of variable loci, mean pairwise differences (π) computed on all sites of variable loci, and inbreeding coefficient (FIS), Tajima’s D and Fu & Li’s D. All Tajima’s D are significant. Maximum (in bold) and minimum (in italics and underlined) values are highlighted for all columns.

| Species | *N* | No. RAD<br>Seq loci | No. variable<br>RADSeq loci<br>(%) | No. SNP (%) | Ho | He | FIS | $\Theta_w$ ($\times 10^{-4}$) | $\pi$ ($\times 10^{-4}$) | Tajima's D | Fu & Li's D |
| --- | --- | --- | --- | --- | --- | --- | --- | --- | --- | --- | --- |
| *Myripristis tiki* | 18 | 45,432 | **28,855** (64) | **46,718 (0.85)** | 0.159 ($\pm 0.021$) | 0.160 ($\pm 0.022$) | 0.025 | 32.54 | 22.23 | -1.211 | -1.586 |
| *Centropyge hotumatua* | 15 | 29,195 | 21,684 (74) | 38,816 (1.09) | 0.156 ($\pm 0.018$) | 0.158 ($\pm 0.016$) | 0.032 | **37.65** | 24.38 | -1.383 | -1.591 |
| *Ostorhinchus chalcus* | 19 | **63,590** | 24,019 (<u>38</u>) | 32,224 (<u>0.42</u>) | 0.165 ($\pm 0.024$) | 0.162 ($\pm 0.021$) | <u>0.008</u> | <u>26.61</u> | <u>18.55</u> | -1.149 | -1.726 |
| *Chaetodon litus* | 15 | 41,439 | 21,905 (53) | 33,422 (0.67) | 0.220 ($\pm 0.029$) | 0.219 ($\pm 0.023$) | 0.022 | 32.09 | 28.86 | -0.395 | -0.500 |
| *Sargocentron wilhelmi* | 16 | 41,234 | 24,207 (59) | 37,861 (0.76) | 0.217 ($\pm 0.028$) | 0.217 ($\pm 0.022$) | 0.026 | 32.36 | 29.19 | -0.381 | -0.220 |
| *Coris debueni* | 14 | 47,949 | 28,594 (60) | 45,870 (0.79) | 0.208 ($\pm 0.027$) | 0.209 ($\pm 0.022$) | 0.029 | 34.35 | 28.94 | -0.624 | -0.747 |
| *Cantherhines rapanui* | 15 | 44,233 | 20,052 (45) | <u>28,239</u> (0.53) | **0.262 ($\pm 0.032$)** | **0.261 ($\pm 0.024$)** | 0.023 | 29.62 | **31.63** | 0.266 | 0.293 |
| *Chrysiptera rapanui* | 18 | <u>25,331</u> | <u>19,246</u> (**76**) | 35,735 (**1.15**) | <u>0.137</u> ($\pm 0.018$) | <u>0.141</u> ($\pm 0.017$) | 0.039 | 37.31 | 22.48 | -1.519 | -2.277 |
| *Chromis randalli* | 13 | 34,893 | 24,673 (71) | 43,845 (1.03) | 0.161 ($\pm 0.019$) | 0.165 ($\pm 0.017$) | **0.043** | 38.81 | 25.47 | -1.378 | -1.814 |

### 3.2 Population genetic statistics

A summary of the principal statistics is presented in **TABLE 2**. Briefly, the observed heterozygosity values varied from 0.137 ± 0.018 (*Chrysiptera rapanui*) to 0.262 ± 0.032 (*Cantherhines rapanui*) and no departures were observed when compared to expected heterozygosity. The FIS values were overall very low for all species, ranging from 0.008 (*Ostorhinchus chalcius*) to 0.043 (*Chromis randalli*). Watterson’s theta values computed on all sites of variable loci ranged from 26.61 × 10^−4^ (*O. chalcius*) to 37.65 × 10^−4^ (*Centropyge hotumatua*) while π computed on all sites of variable loci varied more extensively, from 18.65 × 10^−4^ (*O. chalcius*) to 31.63 × 10^−4^ (*Cantherhines rapanui*). All Tajima’s D values were significant (**TABLE 2**). *Cantherhines rapanui* displayed the only positive value, suggesting a population contraction (bottleneck), while the remaining eight species displayed negative values, suggesting population growth.

We found that none of these population genetic statistics were correlated with other factors such as family, demography, range-size classification, reproductive strategy, or generation time after an MRT analysis. The two-leaf MRT for normalized π, *Θ*_*w*_, Ho, Tajima’s D and Fu & Li’s D explained a high proportion of total variance (67.9%, **FIGURE 1A**) and divided the dataset into two clusters: the first cluster (Cluster 1) was composed of both large range and small range endemics, included all types of reproductive strategies and the lowest π, *Θ*_*w*_, Ho, Tajima’s D and Fu & Li’s D values (*Centropyge hotumatua, Ostorhinchus chalcius, Chrysiptera rapanui, Chromis randalli, Myripristis tiki*). Conversely, the second cluster (Cluster 2) displayed the highest values for all summary statistics (*Sargocentron wilhelmi, Cantherhines rapanui, Chaetodon litus, Coris debueni*). This result was confirmed by visual inspection of the SFS, with species of the first cluster displaying a spectrum fitting a clear expansion while species of the second cluster displayed a spectrum supporting either a bottleneck (*Cantherhines rapanui*) or weak expansion (*Chaetodon litus, Coris debueni* and *Sargocentron wilhelmi*) (**FIGURE 1B**).

**FIGURE 1.**
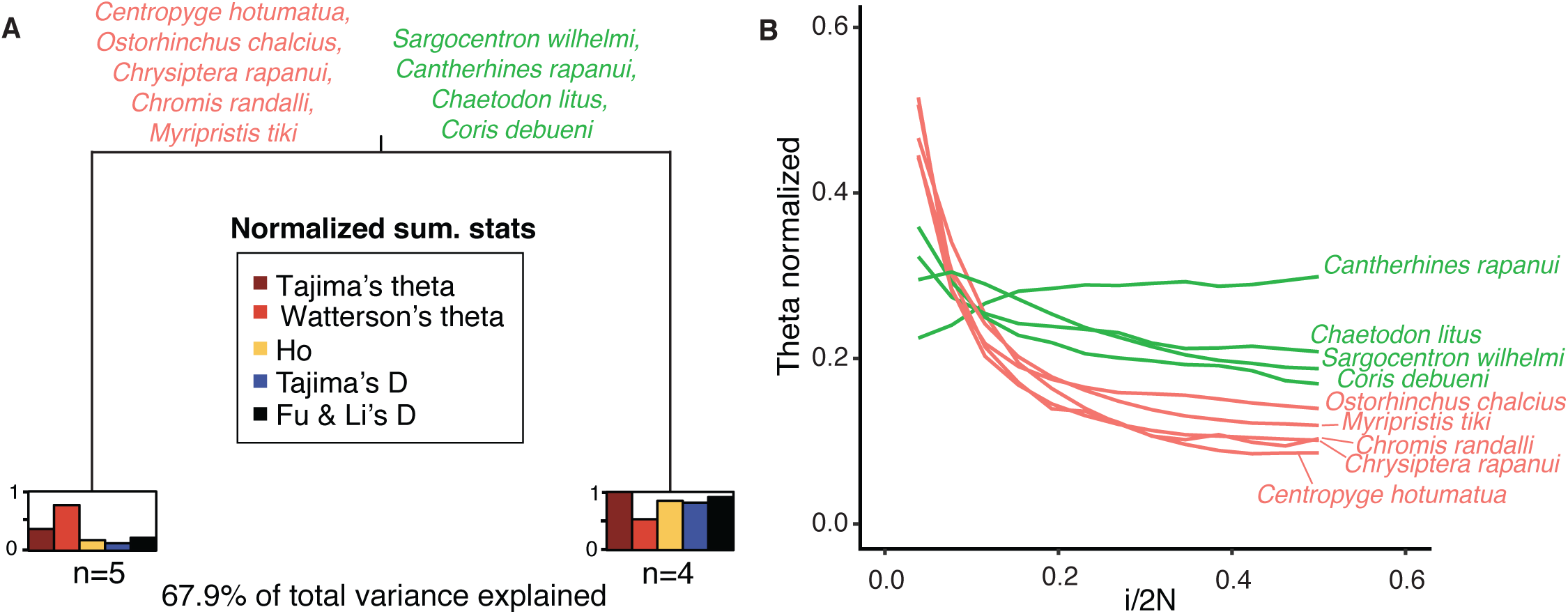
**(A)** Multivariate regression tree (MRT) of normalized summary statistics using as predictor variables range-size classification (large-range endemic and small- range endemic), demographic pattern (expansion, bottleneck), reproductive strategy (pelagic eggs, demersal eggs and mouth brooding), and generation time; **(B)** Aggregated SFS for the nine species being studied. Colors denote the clusters retrieved by the MRT analysis.

### 3.3 Changes in historical effective population size

As both *stairwayplots* and the abc skylines were used to compute changes in effective population sizes, we chose to consider only changes in Ne observed with both methods. We recovered concordant signals with the two methods, with signatures of population expansions for eight out of the nine species and signature of a bottleneck for only one species (**FIGURE 2, 3**). Expansion times retrieved were highly similar between the two methods for most species in Cluster 1: *Centropyge hotumatua* (ca 75,000 ypb); *Chromis randalli* (ca 100,000 ypb); *Chrysiptera rapanui* (ca 50,000 ypb); *Myripristis tiki* (ca 130,000 ypb), but differed for *Ostorhinchus chalcius* (20,000 (stairway) vs 125,000 (abc) ypb). For Cluster 2, abc skylines dated older demographic events for the bottleneck retrieved in *Cantherhines rapanui* (22,000 (stairway) vs 50,000 (abc) ypb) and for the older expansion times retrieved in *Chaetodon litus* (40,000 (stairway) vs 200,000 (abc) ypb), and *Coris debueni* (140,000 (stairway) vs 250,000 (abc) ypb); *Sargocentron wilhelmi* (180,000 (stairway) vs 380,000 (abc) ypb)) (**FIGURE 2, 3**).

**FIGURE 2.**
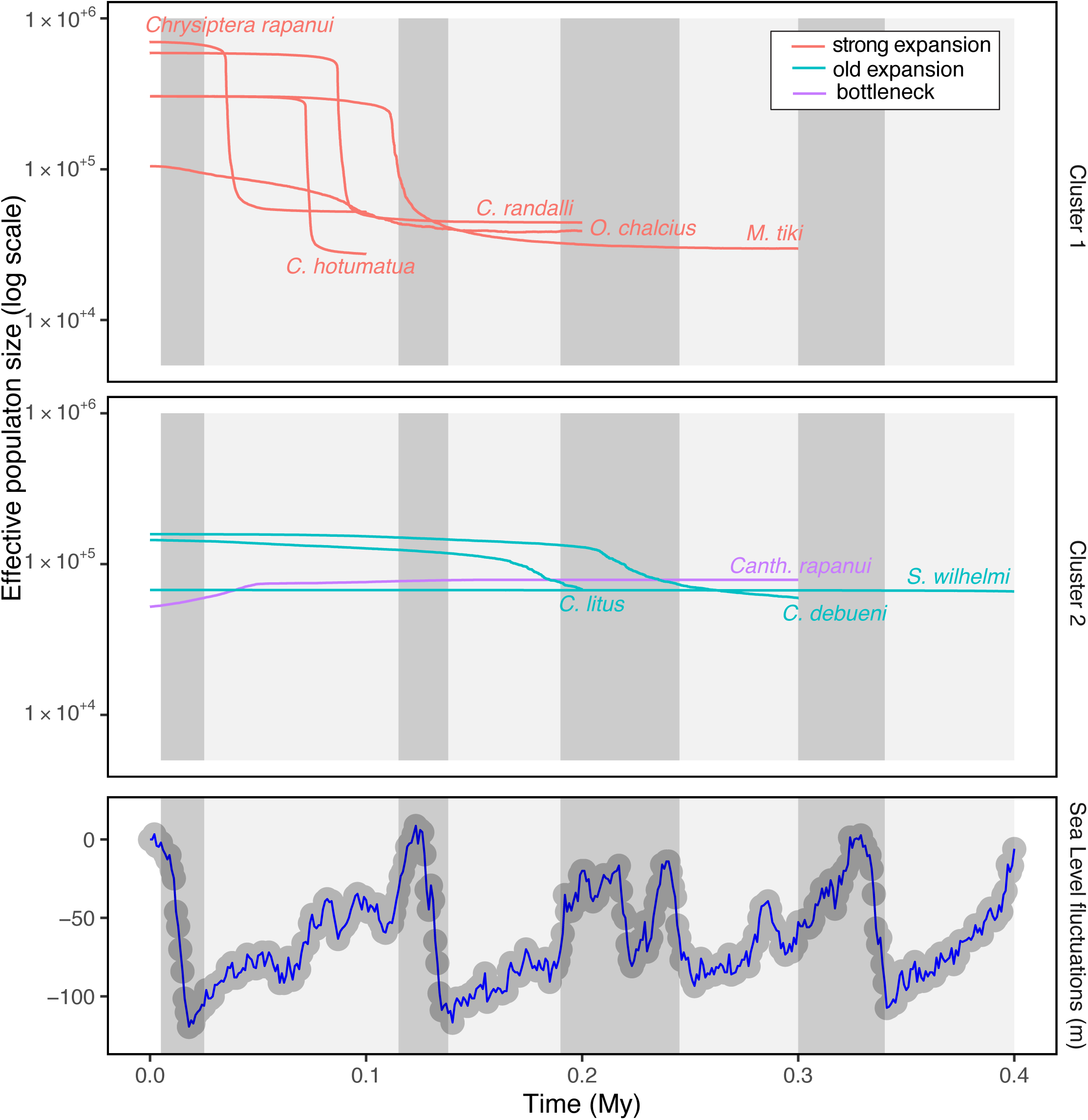
ABC skyline plots representing the variation of the median effective population size through time and sea level fluctuations for the past 400,000 years using data from (Miller et al., 2005). Dark tones indicate inter-glacial periods characterized by sea level rises while grey tones indicate glacial periods with lower sea levels.

**FIGURE 3.**
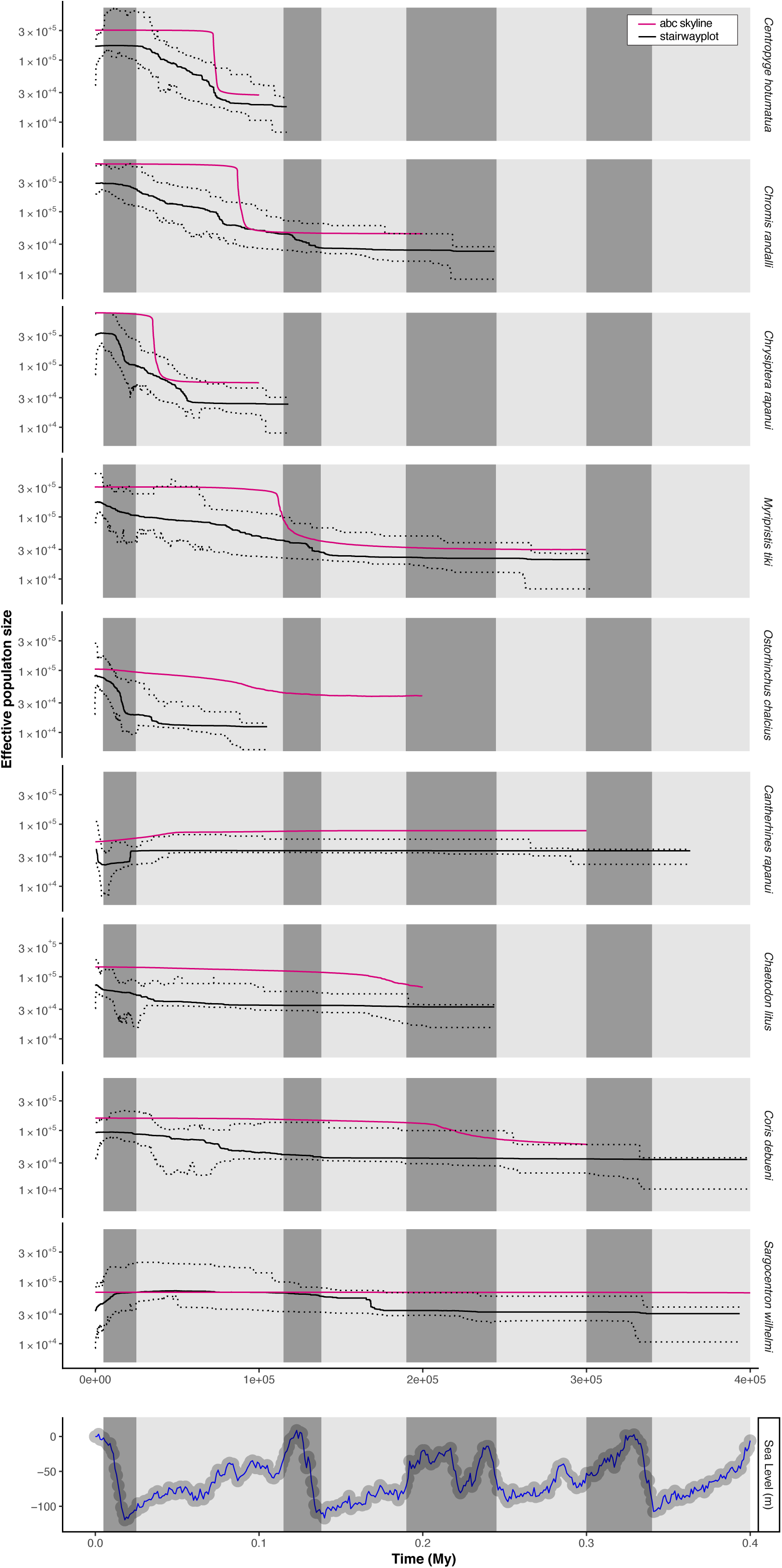
*Stairwayplots* (black) (red) and abc skylines (pink) representing variation in effective population size through time and sea level fluctuations for the past 400,000 years using data from (Miller et al., 2005). Only median Ne is represented for the abc skyline plots while median and 97.5% interval values are represented for the *stairwayplots*. Dark tones indicate inter-glacial periods characterized by sea level rises while grey tones indicate glacial periods with lower sea level.

The hABC analyses allowed us to test the synchronicity of the observed changes in Ne. The estimate for *ζ* was consistent with a concerted expansion for most species, as the median of the *ζ* posterior distribution obtained was 6 (**FIGURE 4**). Cross-validation suggested that the median is the best point estimate for *ζ* (**FIGURE S2**), while the mode did not perform well, with most datasets displaying 1 or 9 no matter the real value of *ζ*. We retrieved a co-expansion time of 154,200 ypb (97.5% CI: 64,687 - 339,057 ypb) and an older mean time (E(τ) = 187,437 ypb, 97.5% CI: 156,140 - 239,756). Restricting the analysis to the species displaying the clear expansions (species of Cluster 1), we found a stronger and younger pulse of coexpansion (τ_s_ = 45,836 ypb, 97.5%, CI: 30,023-87,767 ypb) associated with a younger mean expansion time (E(τ): 82,471 ypb, 97.5% CI: 69,691-107,510 ypb) (**FIGURE 4**). The posterior predictive test highlighted that the estimated model could reproduce the observed data, as shown by the PCA computed on the aSFS of the pseudo-observed and observed data (**FIGURE S3**). The mean TMRCA retrieved with the *stairwayplots* for the nine species was very recent 253,871 ypb (+/- 119,482 ypb) and ranged from 105,232 ypb (*Ostorhinchus chalcius*) to 398,058 ypb (*Sargocentron wilhelmi*) (**FIGURE 1**). Including normalized TMRCAs in the previous MRT analyses resulted in the formation of the same two partitions, with species of Cluster 1 having younger TMRCAs than species of Cluster 2 (65.6% of variance explained, **FIGURE S1**).

**FIGURE 4.**
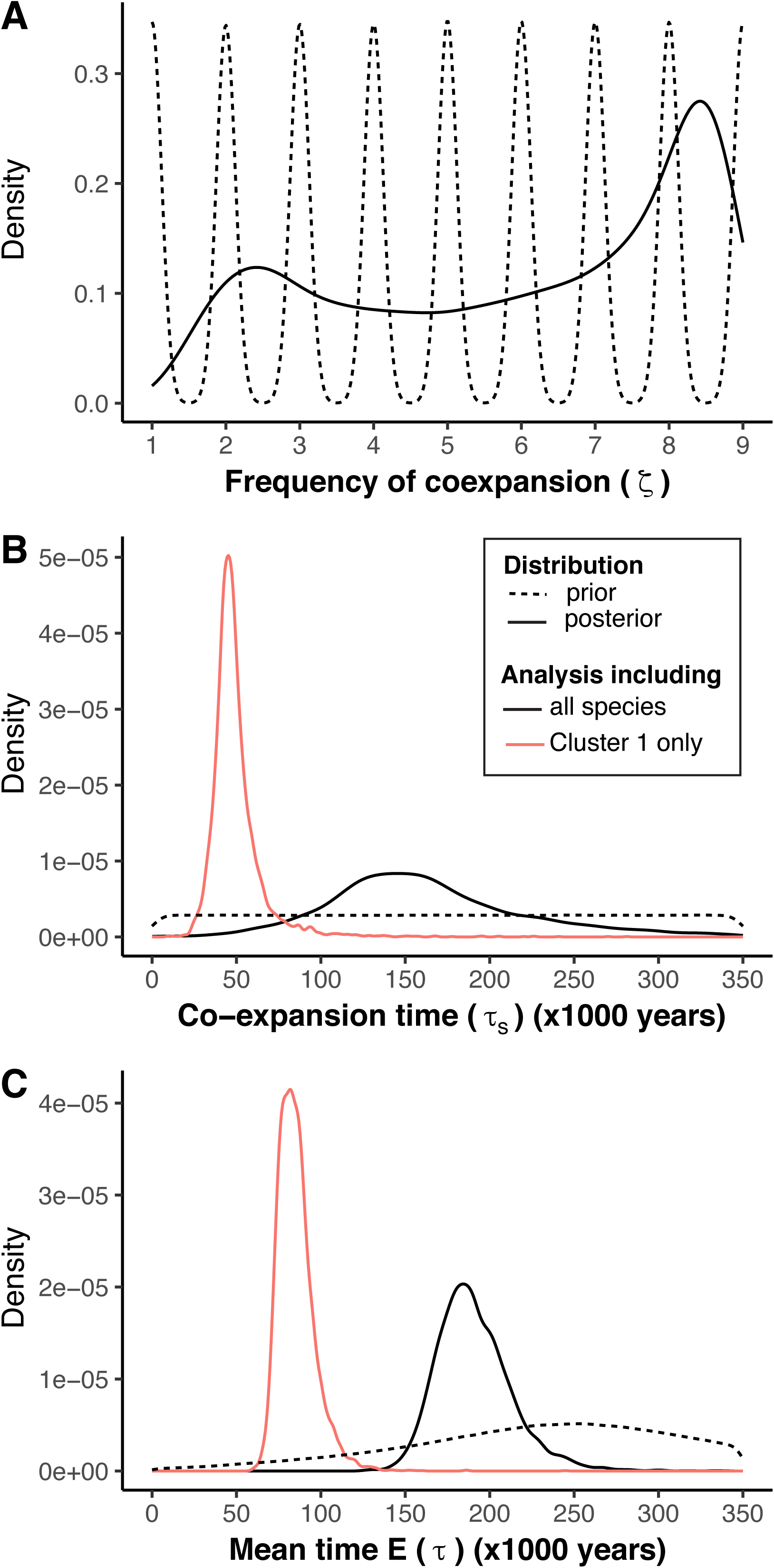
Posterior estimates for the different parameters of the hABC analysis: (A) the frequency of coexpansion (ζ), (B) the co−expansion time (τ), and (C) the total mean expansion time E (τ).

Overall, we recovered three different demographic patterns, concordant with the aSFS profiles and the neutrality tests obtained. (a) The five species of Cluster 1 displayed high magnitude population expansions, with modern Ne being on average 11.8 times higher than the corresponding ancient Ne. (b) Older and weaker expansions were found for two species (*Coris debueni, Chaetodon litus, Sargocentron wilhelmi*), with modern Ne being on average 2.8 times higher than their ancient Ne. (c) Lastly, a bottleneck was found for *Cantherhines rapanui*, with modern Ne 1.6 times lower than the ancient Ne.

### 3.4 Reconstruction of the surface of potential habitat

Reconstruction of the potential present and past habitat during periods of low sea level conditions (−120 m) showed that rise in sea level resulted in a reduction of 38.6% of the potential habitat (**FIGURE 5**). This reduction in potential habitat was accompanied by a reduction in the number of potential seamounts colonized, from 18 to 11 (**FIGURE 5**). It is worth noting that the Apollo seamount was not recovered as a present nor a past habitat despite the fact that the species *Sargocentron wilhelmi* and *Chaetodon litus* have been observed there recently (**TABLE 1**); the resolution of the depth gradient might be the cause of this discrepancy.

**FIGURE 5.**
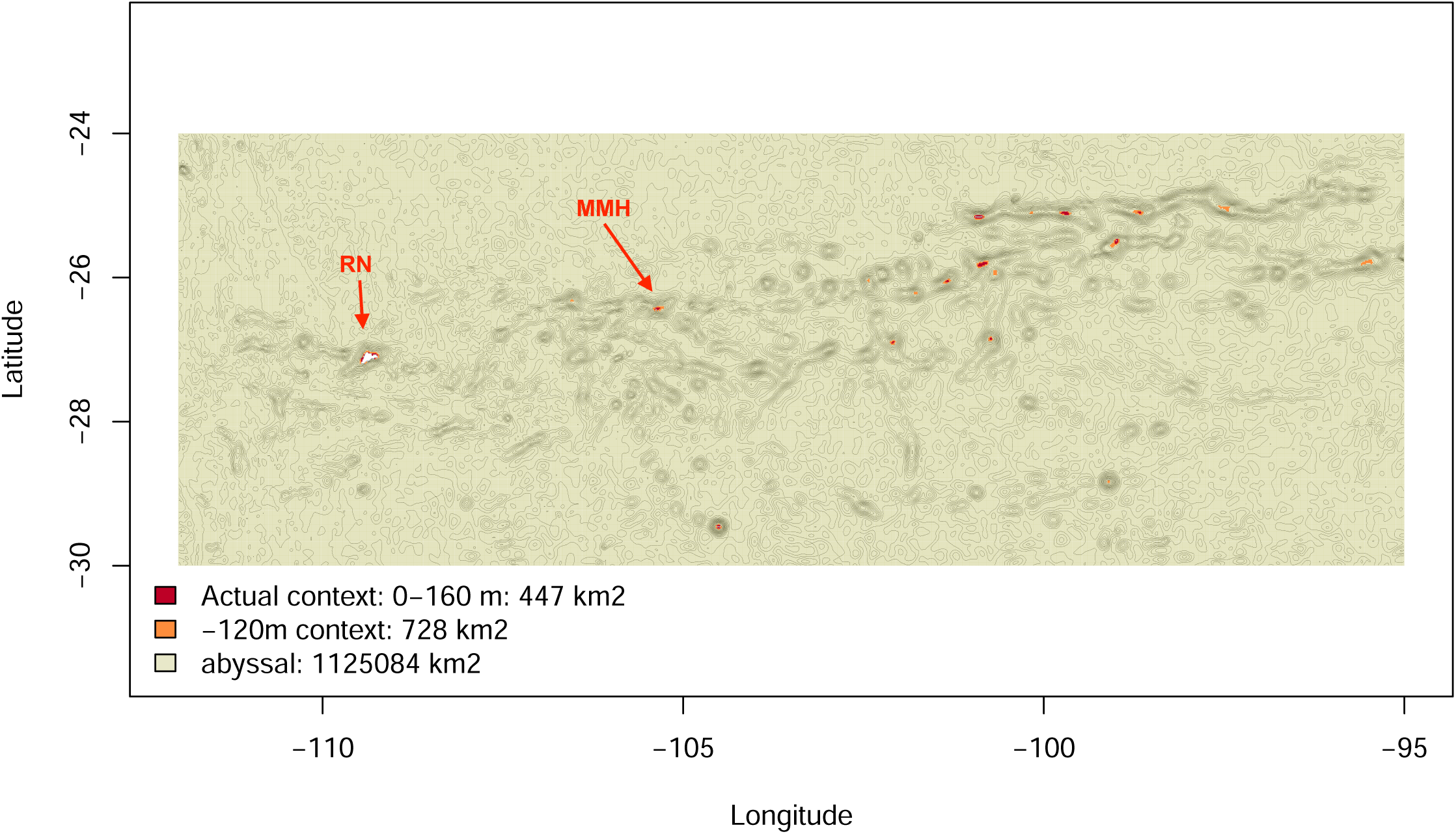
Surface of potential habitat for Rapa Nui endemic reef fish species represented for present conditions (red) and during sea level lowstand conditions with sea level as low as -120m compared to present sea level (orange).

## 4 DISCUSSION

### 4.1 Genetic diversity and demographic changes - identifying common patterns among species

Identifying general patterns in ecosystems is of fundamental importance to understand species responses to environmental changes and underlying eco-evolutionary dynamics. Studies combining multiple species in joint demographic analyses at the community-scale are still scarce in the literature, but they will progressively shed light on common past demographic changes and eventually help to predict future dynamics. Here we harnessed the power of population genomics to uncover the historical demography of one of the most remote coral fish assemblage of the Pacific Ocean. To detect the co-occurrence of similar demographic responses, we first inspected the genetic variability of the nine species and then inferred their demographic history individually through coalescent-based analyses. Overall, we found that Rapa Nui endemic species share a common history, with population expansions dominating the demographic histories of eight out of the nine species studied here. We found signatures of demographic changes that could be dated for most of the species to the last glacial period for both small-range and large-range endemics. We identified one cluster of five species showing a strikingly similar pattern of population expansions (Cluster 1), and a second cluster composed of mildly expanding populations and one species (*Cantherhines rapanui*) experiencing a contraction (**FIGURE 1**,**2**,**3; TABLE 2**). We quantitatively confirmed this result using an hABC model, which found a group of six co-varying species dated at 154,200 ybp (97.5% CI: 64,687 - 339,057 ypb; **FIGURE 4**). The hABC requires, however, a larger number of species to be included in the analysis in order to detect synchronous changes in effective population size (Chan et al. 2014), and our results of cross validation suggest that even with genomic data there is some uncertainty regarding the exact number of co-varying species (**FIGURE S2**).

Combining all of the descriptive and inferential evidences, we therefore ran the hABC analysis on Cluster 1 only to refine the estimation of the timing of co-expansion. A stronger and younger signal of co-variation has been retrieved and dated the coexpansion at 45,836 ypb (97.5%, CI: 30,023-87,767 ypb; **FIGURE 4**), confirming the presence of two clusters among expanding species.

### 4.2 Imprint of sea-level fluctuations on the demographic history of Rapa Nui reef fish species

Sea level oscillations over the past 800,000 years (Augustin et al., 2004; McManus, 2004) and particularly during the Last Glacial Maximum (LGM, from 19,000–20,000 to 26,500 years ago, Clark et al. (2009)) have been hypothesized as greatly affecting coral reef fish populations as coral reef fish species commonly studied are restricted to shallow waters (0-30 m depth, Craig, Eble, & Bowen, 2010; Craig, Eble, Bowen, & Robertson, 2007; Crandall, Frey, Grosberg, & Barber, 2008; Fauvelot, Bernardi, & Planes, 2003; Gaither, Toonen, Robertson, Planes, & Bowen, 2010; Hellberg, 2001; Horne, van Herwerden, Choat, & Robertson, 2008; Klanten, Choat, & Van Herwerden, 2007; Robertson, 2004; Winters, van Herwerden, Choat, & Robertson, 2010). Rapa Nui endemics, however, differ in their ecology compared to organisms of other reefs in the Pacific region, as they seem far less restricted in terms of habitat and depth range. Many of the Rapa Nui endemic species have the particularity of being found from shallow waters to the mesophotic zone, down to at least 160 m in Rapa Nui waters and in nearby seamounts (**TABLE 1**). Considering the network of seamounts surrounding Rapa Nui, low sea levels would have produced periods of maximum reef habitat extension for Rapa Nui endemic species. As such, population expansions observed for most species during the glacial period, a period characterized by climatic cooling and decreased sea level, could reflect the colonization / recolonization history of Rapa Nui from Motu Motiro Hiva or other seamounts of the Rapa Nui archipelago. While the Ancient Archipelago hypothesis of Newman & Foster (1983) has been invalidated at the species level for this island system (Delrieu-Trottin et al., 2019), this hypothesis could explain what is observed at the population level for most Rapa Nui endemic species (i.e. a recolonization from the seamounts rather than from elsewhere in the Pacific).

We would expect to recover a recent bottleneck for all species following sea level rise and corresponding reduction of habitat, a bottleneck correlated with the LGM (Fauvelot et al., 2003; Ludt & Rocha, 2015) was only observed for *Cantherhines rapanui*. This species, together with the apogonid *Ostorhinchus chalcius*, possesses one of the most restricted depth ranges among the species studied here (0-20m, see **TABLE 1**). However, it is important to note that *O. chalcius* is a nocturnal predator and is cryptic during the day. Therefore, it is highly likely that its current depth range is an underestimation that is limited to technical constraints of mesophotic fish surveys. As such, this assumption could explain to some extent why we did not find signal of a bottleneck for this species. In contrast, the restricted depth range of *Cantherines rapanui* could be the reason why the demographic pattern of this species differs from that of the other studied species. More specifically, a restricted depth range would have made this species more vulnerable to sea level fluctuations during the LGM compared to the other Rapa Nui endemics. The ability to thrive across a large depth gradient might have buffered the influence of the LGM on most Rapa Nui endemic populations. Mesophotic reefs usually host different communities of species (Rocha et al., 2018), a change that is attributed to either competition and/or adaptation to such habitats. The low species diversity in Rapa Nui might be one of the factors influencing the presence of Rapa Nui endemics with the capability of inhabiting such large depth gradients.

### 4.3 Community assembly in remote islands – importance of local processes

The common demographic and temporal patterns found for both large-range and small-range endemics also provide insights into the mechanisms that shaped the Rapa Nui ichthyofauna. Community assembly in remote islands is expected to rely on colonization / extinction / recolonization processes. Populations of large-range endemics sampled around Rapa Nui could potentially operate at a regional scale. However, we observed concordant expansion times for species of Cluster 1 (*Centropyge hotumatua, Ostorhinchus chalcius, Chrysiptera rapanui, Chromis randalli, Myripristis tiki*) including two large range endemics, suggesting that population demographic histories have been driven by local historical processes and that neighboring populations have minimally affected the demography of these species.

This is consistent with the isolation of Rapa Nui, which is mirrored by its relatively lower species richness compared to that of other Pacific islands. The low species diversity in Rapa Nui reefs might be due not only to the difficulty of larvae to colonize such remote islands, but also due to the difficulty of establishing a viable population / metapopulation that functions at such local scales. This hypothesis is reinforced by the fact that the two large range endemics studied here are among the most abundant species of Rapa Nui (Friedlander et al., 2013), yet they are rare throughout the rest of their distribution (Randall, 2005). Assessing the amount of gene flow between Rapa Nui populations with those of Pitcairn or Austral Island for these large range endemics could help to test this hypothesis. The predominance of local over regional processes on the population demography of remote endemic species has also been reported for Marquesan endemic reef fish species (Delrieu-Trottin et al., 2017) and for species endemic to the oceanic islands of the Vitória–Trindade Chain (Pinheiro et al., 2017), arguing in favor of potentially common processes for peripheral hotspots of endemism.

### 4.4 Limitations of the demographic inferences

We reconstructed the demographic history of the Rapa Nui fauna following three independent approaches that have complementary strengths and shortcomings in estimating both the direction, the magnitude and the timing of demographic changes through time. The *stairwayplot* is a “model-flexible” (or model-free) approach similar to skyline plot approaches (Drummond, Rambaut, Shapiro, & Pybus, 2005; Drummond & Rambaut, 2007; Heled & Drummond, 2008; Pybus, Rambaut, & Harvey, 2000). Not restricted to a specific demographic model, Ne is free to vary at each coalescent interval, allowing the exploration of a large model space (Liu & Fu, 2015). The large number of Ne parameters may result difficult to estimate when the SNPs panel is relatively small (Lapierre, Lambert, & Achaz, 2017). Moreover, we are specifically interested in the time when a change in Ne occurred, which is only indirectly estimated by the *stairwayplot* as a function of the changes in Ne. On the other hand, the Approximate Bayesian Computation (ABC) framework explicitly modeled three-time parameters while the variation of Ne through time is reconstructed a posteriori (Maisano Delser et al., 2016), having the drawback of approximating the likelihood of the data using summary statistics. For these reasons, *stairwayplot* and the abc skyline are highly complementary and it is important to run both of them to gain a robust insight into species demography. The discrepancy we observed in the timing of change between the two approaches, but not in the trend, it is clearly related to the large variance associated with the estimation process. More genomic data or more species are needed to minimize this variance and the differences observed between the *stairwayplot* and abc skyline results. Here we combined more species using the hABC approach to quantitatively test the synchronicity of the changes in Ne (Chan et al., 2014). The hABC approach is necessarily limited by an a priori choice of the demographic models characterizing the species under investigation. Here we adopted a simple three parameters model because all previous analyses on single species have suggested it could be enough to describe the dynamics of Ne through time. Overall, the addition of more species to these analyses as the power to correctly infer the number of co-varying species increases for larger datasets (Chan et al., 2014).

Finally, the selection of the mutation rates and generation times can greatly impact the timing and the magnitude of the demographic estimates. During the last 20 years, allozymes, microsatellites and Sanger sequence data have mainly been used to investigate the population histories of Pacific shore fishes ; inferences based on SNPs are still scarce (Crane et al., 2018; Maisano Delser et al., 2019). No SNP mutation rate (µ) estimates were available for any of the species of this study. Yet, Brumfield, Beerli, Nickerson, & Edwards, (2003) suggest that SNPs should have relatively low mutation rates, of the order of 10^−8^–10^−9^. For fishes, a large array of mutation rates have been hypothesized. To our knowledge, these range from 2.5 × 10^− 8^ (Kavembe, Kautt, Machado-Schiaffino, & Meyer, 2016; Nunziata & Weisrock, 2018) using a SNP mutation rate calibrated on humans (Excoffier, Dupanloup, Huerta-Sánchez, Sousa, & Foll, 2013b) to as slow as 3.5 × 10^−9^ (Malinsky et al., 2018) using mutation rate estimated in three Lake Malawi cichlids comparing the genotypes of parents and their offspring. The SNP mutation rate we chose (1 × 10^−8^) corresponds to one of the most often used in the literature for fishes (Jacobs et al., 2018; Le Moan et al., 2016; Rougeux et al., 2017; Souissi et al., 2018; Tine et al., 2014) and is in accordance with Brumfield, Beerli, Nickerson, & Edwards, (2003). With this mutation rate, we found relatively high estimates of the effective population sizes (10^5^ to 10^6^); yet a mutation rate of 10^−9^ would have led to even higher estimates, which is less likely considering the fact that the species studied here have very restricted distributions. In the same manner, our estimates of the expansion times found with this mutation rate are in agreement with previous studies of the demographic history of Pacific reef fishes based on mitochondrial sequence data (Delrieu-Trottin et al., 2017; Dibattista, Rocha, Craig, Feldheim, & Bowen, 2012; Gaither et al., 2010), which have recovered population expansions during the last 400,000 years. Generation times can also strongly bias estimates, resulting for instance in synchronous population fluctuations among species with similar generation times. Alternatively, applying erroneous generation times to species with similar demographic trajectories can lead to the spurious inference of asynchronous demographic changes. In order to use generation time estimates that are as accurate as possible, we selected generation times of close relatives and of similar sized species from the literature for most species (**TABLE 1**). The fact that the most recent demographic change was found for the species with the greatest generation time (*Cantherines rapanui*, 3 years) and the Cluster of coexpanding species (Cluster 1) comprises five species with three different generations times together suggest that the synchronous demographic expansions inferred here are not spurious and results from community dynamics associated with eustatic changes. This being said, estimates of generation times and mutation rates for more species of fishes will strengthen future demographic analyses.

## 5 CONCLUSIONS

We elucidate for the first time the demographic history of the Rapa Nui endemic reef fish community using genomic data. We show that the Rapa Nui reef fish endemic community shares a common history, with expansions occurring during the last glacial period for the two different types of endemic species. Local processes based on the seamount system around Rapa Nui have played a major role in the foundation and persistence of this endemic reef fish community through dynamic colonization / extinction / recolonization events throughout the Rapa Nui Archipelago. Altogether, our results reflect the importance of working at the community level, i.e. exploring multiple species to uncover general trends. Finally, our results provide insights on the mechanisms by which peripheral endemism has originated in coral reef fishes.

## ACKNOWLEDGEMENTS

We thank Rebeca Tepano, Nina, Taveke Olivares Rapu, Liza Garrido Toleado (SERNAPESCA), Ludovic Tuki (Mesa del Mar, Te Mau O Te Vaikava O Rapa Nui), and the people of the Rapa Nui Island for their kind and generous support. We thank Luiz A. Rocha for kindly sharing photographs and information related to maximum depth of several Rapa Nui endemic species. We are grateful to the Genotoul bioinformatics platform Toulouse Midi-Pyrenees (Bioinfo Genotoul; http://bioinfo.genotoul.fr/) for providing computing resources. We are obliged to three anonymous reviewers for their constructive and helpful advices. This work was supported by the FONDECYT Postdoctorado fellowship N°3160692 to E. Delrieu-Trottin and by the ATM grant 2016 from the Muséum National d’Histoire Naturelle to Stefano Mona. E.C.Giles thanks the scholarship BECA CONICYT-PCHA/Doctorado Nacional Chile/2019 Folio No. 21190878. P. Saenz-Agudelo was supported by FONDECYT grant N°1190710.

## ETHICS

All applicable institutional guidelines for the care and use of animals were followed. Specimens were collected under permit No. 724, 8 March 2016 obtained from the Chilean Subsecretary of Fishing. The Universidad Austral de Chile Ethical Care Committee and Biosecurity Protocol approved our use and handling of animals.

## AUTHOR CONTRIBUTIONS

E.D.-T. and P.S.-A. conceived the study; E.D.-T., P.S.-A. and S.M. acquired the funding; E.D.-T., E.C.G., V.N, C.R.E. and P.S.-A., collected the field data; E.D.-T., P.C.- B. and A.S. produced the data; E.D.-T., N.H., S.M, and P.S.-A. analyzed the data; E.D.-T., N.H., E.C.G., S.M, and P.S.-A wrote the manuscript. All authors gave final approval for publication.

## DATA ACCESSIBILITY

All Fastq sequence files are available from the GenBank at the National Center for Biotechnology Information short-read archive database (BioProject accession number: PRJNA622670).

## Supplementary Material

**FIGURE S1** Multivariate regression tree (MRT) of normalized summary statistics using as predictor variables range-size classification (large-range endemic and small-range endemic), demographic pattern (expansion, bottleneck), reproductive strategy (pelagic eggs, demersal eggs and mouth brooding), and generation time.

**FIGURE S2** Cross-validation for ζ after 1,000 simulations from prior distributions. X- axe: simulated ζ. Y-axe: estimated ζ through the same ABC procedure implemented for the real data.

**FIGURE S3** PCA computed on the posterior predictive distribution of the aSFS after 50,000 simulations. Black dot: pseudo-observed (simulated) datasets; red dot: observed data. The first two axes displayed here represent 62.98 % of the total variance.

## REFERENCES

Allen, G. R. (2008). Conservation hotspots of biodiversity and endemism for Indo-Pacific coral reef fishes. Aquatic Conservation: Marine and Freshwater Ecosystems, 18(5), 541–556. doi:10.1002/aqc.880

Allen, G. R., & Erdmann, M. (2012). Reef fishes of the East Indies. …I--III. Tropical Reef …, 1292. Retrieved from http://www.uhpress.hawaii.edu/p-8881-9780987260000.aspx

Augustin, L., Barbante, C., Barnes, P. R. F., Barnola, J. M., Bigler, M., Castellano, E., … EPICA community members. (2004). Eight glacial cycles from an Antarctic ice core. Nature. doi:10.1038/nature02599

Avise, J. C. (2009). Phylogeography: retrospect and prospect. Journal of Biogeography, 36(1), 3–15.

Bard, E., Hamelin, B., Arnold, M., Montaggioni, L., Cabioch, G., Faure, G., & Rougerie, F. (1996). Deglacial sea-level record from Tahiti corals and the timing of global meltwater discharge. Nature, 382(6588), 241–244. doi:10.1038/382241a0

Beaumont, M. A., Zhang, W., & Balding, D. J. (2002). Approximate Bayesian computation in population genetics. Genetics.

Bellwood, D. R., & Wainwright, P. C. (2002). The History and Biogeography of Fishes on Coral Reefs. In Coral Reef Fishes (pp. 5–32). doi:10.1016/B978-012615185-5/50003-7

Bowen, B. W., Muss, A., Rocha, L. A., & Grant, W. S. (2006). Shallow mtDNA coalescence in Atlantic pygmy angelfishes (genus Centropyge) indicates a recent invasion from the Indian Ocean. Journal of Heredity, 97(1), 1–12. doi:10.1093/jhered/esj006

Breiman, L., Friedman, J. H., Olshen, R. A., & Stone, C. J. (1984). Classification and regression trees. Belmont, CA: Wadsworth. International Group, 432, 151–166.

Brumfield, R. T., Beerli, P., Nickerson, D. A., & Edwards, S. V. (2003). The utility of single nucleotide polymorphisms in inferences of population history. Trends in Ecology and Evolution, 18(5), 249–256. doi:10.1016/S0169-5347(03)00018-1

Burbrink, F. T., Chan, Y. L., Myers, E. A., Ruane, S., Smith, B. T., & Hickerson, M. J. (2016). Asynchronous demographic responses to Pleistocene climate change in Eastern Nearctic vertebrates. Ecology Letters, 19(12), 1457–1467. doi:10.1111/ele.12695

Catchen, J., Hohenlohe, P. A., Bassham, S., Amores, A., & Cresko, W. A. (2013). Stacks: An analysis tool set for population genomics. Molecular Ecology, 22(11), 3124–3140. doi:10.1111/mec.12354

Catchen, J. M., Amores, A., Hohenlohe, P., Cresko, W., & Postlethwait, J. H. (2011). Stacks : Building and Genotyping Loci De Novo From Short-Read Sequences. G3 Genes|Genomes|Genetics, 1(3), 171–182. doi:10.1534/g3.111.000240

Cea, A. (2016). Ika Rapa Nui. Rapa Nui Press.

Chan, Y. L., Schanzenbach, D., & Hickerson, M. J. (2014). Detecting concerted demographic response across community assemblages using hierarchical approximate Bayesian computation. Molecular Biology and Evolution, 31(9), 2501–2515. doi:10.1093/molbev/msu187

Clark, P. U., Dyke, A. S., Shakun, J. D., Carlson, A. E., Clark, J., Wohlfarth, B., … McCabe, A. M. (2009). The last glacial maximum. Science, 325(5941), 710–714.

Clouard, V., & Bonneville, A. (2005). Ages of seamounts, islands, and plateaus on the Pacific plate. Special Paper 388: Plates, Plumes and Paradigms, 388, 71–90. doi:10.1130/0-8137-2388-4.71

Craig, M. T., Eble, J. A., & Bowen, B. W. (2010). Origins, ages and population histories: Comparative phylogeography of endemic Hawaiian butterflyfishes (genus Chaetodon). Journal of Biogeography, 37(11), 2125–2136. doi:10.1111/j.1365-2699.2010.02358.x

Craig, M. T., Eble, J. A., Bowen, B. W., & Robertson, D. R. (2007a). High genetic connectivity across the Indian and Pacific Oceans in the reef fish Myripristis berndti (Holocentridae). Marine Ecology Progress Series, 334, 245–254. doi:10.3354/meps334245

Craig, M. T., Eble, J. A., Bowen, B. W., & Robertson, D. R. (2007b). High genetic connectivity across the Indian and Pacific Oceans in the reef fish Myripristis berndti (Holocentridae). Marine Ecology Progress Series, 334, 245–254. doi:10.3354/meps334245

Crandall, E. D., Frey, M. A., Grosberg, R. K., & Barber, P. H. (2008). Contrasting demographic history and phylogeographical patterns in two Indo-Pacific gastropods. Molecular Ecology, 17(2), 611–626. doi:10.1111/j.1365-294X.2007.03600.x

Crane, N. L., Tariel, J., Caselle, J. E., Friedlander, A. M., Ross Robertson, D., & Bernardi, G. (2018). Clipperton Atoll as a model to study small marine populations: Endemism and the genomic consequences of small population size. PLoS ONE. doi:10.1371/journal.pone.0198901

Cutler, K. B., Edwards, R. L., Taylor, F. W., Cheng, H., Adkins, J., Gallup, C. D., … Bloom, A. L. (2003). Rapid sea-level fall and deep-ocean temperature change since the last interglacial period. Earth and Planetary Science Letters, 206(3–4), 253–271.

De’ath, G. (2002). Multivariate Regression Tree: A New Technique for Modeling Species–Environment Relationships. Ecology, 83(4), 1105–1117. doi:10.1890/0012-9658(2002)083[1105:MRTANT]2.0.CO;2

Delrieu-Trottin, E., Brosseau-Acquaviva, L., Mona, S., Neglia, V., Giles, E. C., Rapu-Edmunds, C., & Saenz-Agudelo, P. (2019). Understanding the origin of the most isolated endemic reef fish fauna of the Indo-Pacific: Coral reef fishes of Rapa Nui. Journal of Biogeography, 46(January), 723–733. doi:10.1111/jbi.13531

Delrieu-Trottin, E., Maynard, J., & Planes, S. (2014). Endemic and widespread coral reef fishes have similar mitochondrial genetic diversity. Proceedings of the Royal Society B: Biological Sciences, 281(1797). doi:10.1098/rspb.2014.1068

Delrieu-Trottin, E., Mona, S., Maynard, J., Neglia, V., Veuille, M., & Planes, S. (2017). Population expansions dominate demographic histories of endemic and widespread Pacific reef fishes. Scientific Reports, 7(January), 40519. doi:10.1038/srep40519

Delrieu-Trottin, E., Williams, J. T., Bacchet, P., Kulbicki, M., Mourier, J., Galzin, R., … Planes, S. (2015). Shore fishes of the Marquesas Islands, an updated checklist with new records and new percentage of endemic species. Check List, 11(5). doi:10.15560/11.5.1758

Dibattista, J. D., Rocha, L. A., Craig, M. T., Feldheim, K. A., & Bowen, B. W. (2012). Phylogeography of two closely related indo-pacific butterflyfishes reveals divergent evolutionary histories and discordant results from mtDNA and microsatellites. Journal of Heredity, 103(5), 617–629. doi:10.1093/jhered/ess056

Drummond, A J, Rambaut, A., Shapiro, B., & Pybus, O. G. (2005). Bayesian coalescent inference of past population dynamics from molecular sequences. Molecular Biology and Evolution, 22(5), 1185–1192. doi:10.1093/molbev/msi103

Drummond, Alexei J, & Rambaut, A. (2007). BEAST: Bayesian evolutionary analysis by sampling trees. BMC Evolutionary Biology, 7(1), 214. doi:10.1186/1471-2148-7-214

Easton, E. E., Sellanes, J., Berkenpas, E., Gaymer, C. F., Morales, N., & Gorny, M. (2017). Diversity of deep-sea fi shes of the Easter Island Ecoregion. Deep-Sea Research Part II, 137(December 2016), 78–88. doi:10.1016/j.dsr2.2016.12.006

Eschmeyer, W. N., Fricke, R., Fong, J. D., & Polack, D. A. (2010). Marine fish diversity: History of knowledge and discovery (Pisces). Zootaxa, 2525(2525), 19–50. Retrieved from http://www.mapress.com/zootaxa/

Excoffier, L., Dupanloup, I., Huerta-Sánchez, E., Sousa, V. C., & Foll, M. (2013a). Robust Demographic Inference from Genomic and SNP Data. PLoS Genetics, 9(10). doi:10.1371/journal.pgen.1003905

Excoffier, L., Dupanloup, I., Huerta-Sánchez, E., Sousa, V. C., & Foll, M. (2013b). Robust Demographic Inference from Genomic and SNP Data. PLoS Genetics. doi:10.1371/journal.pgen.1003905

Fauvelot, C., Bernardi, G., & Planes, S. (2003). Reductions in the mitochondrial DNA diversity of coral reef fish provide evidence of population bottlenecks resulting from Holocene sea-level change. Evolution; International Journal of Organic Evolution, 57(7), 1571–1583. doi:10.1111/j.0014-3820.2003.tb00365.x

Friedlander, A. M., Ballesteros, E., Beets, J., Berkenpas, E., Gaymer, C. F., Gorny, M., & Sala, E. (2013). Effects of isolation and fishing on the marine ecosystems of Easter Island and Salas y Gómez, Chile. Aquatic Conservation: Marine and Freshwater Ecosystems, 23(4), 515–531. doi:10.1002/aqc.2333

Froese, R., & Binohlan, C. (2000). Empirical relationships to estimate asymptotic length, length at first maturity and length at maximum yield per recruit in fishes, with a simple method to evaluate length frequency data. Journal of Fish Biology, 56(4), 758–773. doi:10.1111/j.1095-8649.2000.tb00870.x

Gaboriau, T., Leprieur, F., Mouillot, D., & Hubert, N. (2018). Influence of the geography of speciation on current patterns of coral reef fish biodiversity across the Indo-Pacific. Ecography, 41(8), 1295–1306. doi:10.1111/ecog.02589

Gaither, M. R., Toonen, R. J., Robertson, D. R., Planes, S., & Bowen, B. W. (2010). Genetic evaluation of marine biogeographical barriers: Perspectives from two widespread Indo-Pacific snappers (Lutjanus kasmira and Lutjanus fulvus). Journal of Biogeography, 37(1), 133–147. doi:10.1111/j.1365-2699.2009.02188.x

Gajdzik, L., Bernardi, G., Lepoint, G., & Frédérich, B. (2018). Genetic diversity mirrors trophic ecology in coral reef fish feeding guilds. Molecular Ecology, 27(24), 5004–5018. doi:10.1111/mec.14936

Heled, J., & Drummond, A. J. (2008). Bayesian inference of population size history from multiple loci. BMC Evolutionary Biology, 8(1), 289. doi:10.1186/1471-2148-8-289

Hellberg, M. E. (2001). Climate-Driven Range Expansion and Morphological Evolution in a Marine Gastropod. Science, 292(5522), 1707–1710. doi:10.1126/science.1060102

Hewitt, G. (2003). Ice ages: their impact on species distributions and evolution. In Evolution on Planet Earth (eds. L.J. Rothschild & A.M Lister) (pp. 339–361). Academic Press.

Hewitt, G. M. (2001). Speciation, hybrid zones and phylogeography--or seeing genes in space and time. Molecular Ecology, 10(3), 537–549. doi:10.1046/j.1365-294x.2001.01202.x

Hewitt, G. M. (2004). The structure of biodiversity – insights from molecular phylogeography. Frontiers in Zoology, 1(1), 4. doi:10.1186/1742-9994-1-4

Hickerson, M. J., Carstens, B. C., Cavender-Bares, J., Crandall, K. A., Graham, C. H., Johnson, J. B., … Yoder, A. D. (2010). Phylogeography’s past, present, and future: 10 years after. Molecular Phylogenetics and Evolution, 54(1), 291–301.

Horne, J. B., van Herwerden, L., Choat, J. H., & Robertson, D. R. (2008). High population connectivity across the Indo-Pacific: Congruent lack of phylogeographic structure in three reef fish congeners. Molecular Phylogenetics and Evolution, 49(2), 629–638. doi:10.1016/j.ympev.2008.08.023

Hughes, T. P., Bellwood, D. R., & Connolly, S. R. (2002). Biodiversity hotspots, centres of endemicity, and the conservation of coral reefs. Ecology Letters, 5(6), 775–784. doi:10.1046/j.1461-0248.2002.00383.x

Jacobs, A., Hughes, M., Robinson, P., Adams, C., & Elmer, K. (2018). The Genetic Architecture Underlying the Evolution of a Rare Piscivorous Life History Form in Brown Trout after Secondary Contact and Strong Introgression. Genes, 9(6), 280. doi:10.3390/genes9060280

Jombart, T. (2008). Adegenet: A R package for the multivariate analysis of genetic markers. Bioinformatics, 24(11), 1403–1405. doi:10.1093/bioinformatics/btn129

Jombart, T., & Ahmed, I. (2011). adegenet 1.3-1: New tools for the analysis of genome-wide SNP data. Bioinformatics, 27(21), 3070–3071. doi:10.1093/bioinformatics/btr521

Jump, A. S., & Penuelas, J. (2005). Running to stand still: adaptation and the response of plants to rapid climate change. Ecology Letters, 8(9), 1010–1020.

Kavembe, G. D., Kautt, A. F., Machado-Schiaffino, G., & Meyer, A. (2016). Ecomorphological differentiation in Lake Magadi tilapia, an extremophile cichlid fish living in hot, alkaline and hypersaline lakes in East Africa. Molecular Ecology, 25(7), 1610–1625. doi:10.1111/mec.13461

Klanten, O. S., Choat, J. H., & Van Herwerden, L. (2007). Extreme genetic diversity and temporal rather than spatial partitioning in a widely distributed coral reef fish. Marine Biology, 150(4), 659–670. doi:10.1007/s00227-006-0372-7

Knaus, B. J., & Grünwald, N. J. (2017). vcfr: a package to manipulate and visualize variant call format data in R. Molecular Ecology Resources, 17(1), 44–53. doi:10.1111/1755-0998.12549

Lapierre, M., Lambert, A., & Achaz, G. (2017). Accuracy of Demographic Inferences from the Site Frequency Spectrum : The Case of the, 206(May), 439–449. doi:10.1534/genetics.116.192708/-/DC1.1

Le Moan, A., Gagnaire, P. A., & Bonhomme, F. (2016). Parallel genetic divergence among coastal-marine ecotype pairs of European anchovy explained by differential introgression after secondary contact. Molecular Ecology. doi:10.1111/mec.13627

Lessa, E. P., Cook, J. A., & Patton, J. L. (2003). Genetic footprints of demographic expansion in North America, but not Amazonia, during the Late Quaternary. Proceedings of the National Academy of Sciences, 100(18), 10331–10334.

Liu, X., & Fu, Y. X. (2015). Exploring population size changes using SNP frequency spectra. Nature Genetics, 47(5), 555–559. doi:10.1038/ng.3254

Ludt, W. B., & Rocha, L. A. (2015). Shifting seas: The impacts of Pleistocene sea-level fluctuations on the evolution of tropical marine taxa. Journal of Biogeography, 42(1), 25–38. doi:10.1111/jbi.12416

Maisano Delser, P., Corrigan, S., Duckett, D., Suwalski, A., Veuille, M., Planes, S., … Mona, S. (2019). Demographic inferences after a range expansion can be biased: the test case of the blacktip reef shark (Carcharhinus melanopterus). Heredity. doi:10.1038/s41437-018-0164-0

Maisano Delser, P., Corrigan, S., Hale, M., Li, C., Veuille, M., Planes, S., … Mona, S. (2016). Population genomics of C. melanopterus using target gene capture data: demographic inferences and conservation perspectives. Scientific Reports, 6(1), 33753. doi:10.1038/srep33753

Malinsky, M., Svardal, H., Tyers, A. M., Miska, E. A., Genner, M. J., Turner, G. F., & Durbin, R. (2018). Whole-genome sequences of Malawi cichlids reveal multiple radiations interconnected by gene flow. Nature Ecology and Evolution, 2(12), 1940–1955. doi:10.1038/s41559-018-0717-x

Mastretta-Yanes, A., Arrigo, N., Alvarez, N., Jorgensen, T. H., Piñero, D., & Emerson, B. C. (2015). Restriction site-associated DNA sequencing, genotyping error estimation and de novo assembly optimization for population genetic inference. Molecular Ecology Resources, 15(1), 28–41. doi:10.1111/1755-0998.12291

McManus, J. F. (2004). A great grand-daddy of ice cores. Nature. doi:10.1038/429611a

Miller, K. G., Miller, K. G., Kominz, M. A., Browning, J. V, Wright, J. D., Mountain, G. S., … Pekar, S. F. (2005). The Phanerozoic Record of Global Sea-Level Change. Science, 310, 1293–1298. doi:10.1126/science.1116412

Newman, W., & Foster, B. (1983). The Rapanuian faunal district (Easter and Sala y Gomez): in search of ancient archipelagos. Bulletin of Marine Science, 33(3), 633–644.

Nunziata, S. O., & Weisrock, D. W. (2018). Estimation of contemporary effective population size and population declines using RAD sequence data. Heredity, 120(3), 196–207. doi:10.1038/s41437-017-0037-y

Oksanen, J., Blanchet, F. G., Kindt, R., Legendre, P., Minchin, P. R., O’Hara, R. B., … Wagner, H. (2012). vegan: Community Ecology Package. R package version 2.1-20/r2309. Www.R-Project.Org., 1–255. Retrieved from http://r-forge.r-project.org/projects/vegan/

Pahnke, K., Zahn, R., Elderfield, H., & Schulz, M. (2003). 340,000-year centennial-scale marine record of Southern Hemisphere climatic oscillation. Science, 301(5635), 948–952.

Pante, E., & Simon-Bouhet, B. (2013). marmap: A Package for Importing, Plotting and Analyzing Bathymetric and Topographic Data in R. PLOS ONE, 8(9), 1–4. doi:10.1371/journal.pone.0073051

Paradis, E. (2010). pegas: an {R} package for population genetics with an integrated--modular approach. Bioinformatics, 26, 419–420.

Peterson, B. K., Weber, J. N., Kay, E. H., Fisher, H. S., & Hoekstra, H. E. (2012). Double digest RADseq: An inexpensive method for de novo SNP discovery and genotyping in model and non-model species. PLoS ONE, 7(5). doi:10.1371/journal.pone.0037135

Petit, R. J., Aguinagalde, I., de Beaulieu, J.-L., Bittkau, C., Brewer, S., Cheddadi, R., … others. (2003). Glacial refugia: hotspots but not melting pots of genetic diversity. Science, 300(5625), 1563–1565.

Pfeifer, B., Wittelsbürger, U., Ramos-Onsins, S. E., & Lercher, M. J. (2014). PopGenome: An efficient swiss army knife for population genomic analyses in R. Molecular Biology and Evolution, 31(7), 1929–1936. doi:10.1093/molbev/msu136

Pinheiro, H. T., Bernardi, G., Simon, T., Joyeux, J.-C., Macieira, R. M., Gasparini, J. L., … Rocha, L. A. (2017). Island biogeography of marine organisms. Nature, 549, 82. Retrieved from http://dx.doi.org/10.1038/nature23680

Pybus, O. G., Rambaut, A., & Harvey, P. H. (2000). An integrated framework for the inference of viral population history from reconstructed genealogies. Genetics, 155(3), 1429–1437. doi:10.1073/pnas.88.5.1597

R Core Team. (2017). R: A Language and Environment for Statistical Computing. R Foundation for Statistical Computing, Vienna, Austria. doi: http://www.R-project.org/

Randall, J. E. (1998). Review Article Zoogeography of Shore Fishes of the lndo-Pacific Region. ZOOLOGICAL STUDIES, 37, 227–268. Retrieved from http://www.sinica.edu.tw/zool/zoolstud/Journals/37.4/227.pdf

Randall, J. E. (2005). Reef and Shore Fishes of the South Pacific : New Caledonia to Tahiti and the Pitcairn Islands (Vol. 1). University of Hawai’i Press Honolulu.

Randall, J. E. (2007). Reef fishes of Hawaii. University of Hawai’i Press Honolulu. Randall, J. E., & Cea, A. (2011). Shore Fishes of Easter Island. University of Hawai’I Press. Retrieved from http://books.google.com/books?hl=en&lr=&id=iXtCTSRTcb8C&pgis=1

Ray, J. S., Mahoney, J. J., Duncan, R. A., Ray, J., Wessel, P., & Naar, D. F. (2012). Chronology and geochemistry of lavas from the Nazca Ridge and Easter Seamount Chain: An ∼30 myr hotspot record. Journal of Petrology, 53(7), 1417–1448. doi:10.1093/petrology/egs021

Reid, B. N., Naro-Maciel, E., Hahn, A. T., FitzSimmons, N. N., & Gehara, M. (2019). Geography best explains global patterns of genetic diversity and postglacial co-expansion in marine turtles. Molecular Ecology, 28(14), 3358–3370. doi:10.1111/mec.15165

Robertson, D. R. (2004). High genetic diversities and complex genetic structure in an Indo-Paci c tropical reef sh (Chlorurus sordidus): evidence of an unstable evolutionary past? Marine Biology, 757–767. doi:10.1007/s00227-003-1224-3

Rocha, L. A., Pinheiro, H. T., Shepherd, B., Papastamatiou, Y. P., Luiz, O. J., Pyle, R. L., & Bongaerts, P. (2018). Mesophotic coral ecosystems are threatened and ecologically distinct from shallow water reefs. Science, 361(6399), 281–284. doi:10.1126/science.aaq1614

Rougeux, C., Bernatchez, L., & Gagnaire, P. A. (2017). Modeling the multiple facets of speciation-with-gene-flow toward inferring the divergence history of lake whitefish species pairs (Coregonus clupeaformis). Genome Biology and Evolution, 9(8), 2057–2074. doi:10.1093/gbe/evx150

Souissi, A., Bonhomme, F., Manchado, M., Bahri-Sfar, L., & Gagnaire, P. A. (2018). Genomic and geographic footprints of differential introgression between two divergent fish species (Solea spp.). Heredity. doi:10.1038/s41437-018-0079-9

Therneau, T. M., Oksanen, B. R., Oksanen, J., Atkinson, B., & De’ath, G. (2014). mvpart: Multivariate partitioning. Retrieved from https://cran.r-project.org/package=mvpart

Tine, M., Kuhl, H., Gagnaire, P.-A., Louro, B., Desmarais, E., Martins, R. S. T., … Reinhardt, R. (2014). European sea bass genome and its variation provide insights into adaptation to euryhalinity and speciation. Nature Communications, 5(May), 5770. doi:10.1038/ncomms6770

Vrba, E. S. (1992). Mammals as a key to evolutionary theory. Journal of Mammalogy, 73(1), 1–28.

Wakeley, J. (2009). Coalescent theory: an introduction. (ed. Greenwood Village, Colorado: Roberts & Company Publishers)

Wakeley, J. (2010). Natural Selection and Coalescent Theory. Evolution since Darwin: The First 150 Years, 119–149. doi:10.1016/B978-012323448-3/50010-6

Wickham, H. (2009). ggplot2: Elegant Graphics for Data Analysis. Springer-Verlag New York. Retrieved from http://ggplot2.org

Winters, K. L., van Herwerden, L., Choat, J. H., & Robertson, D. R. (2010). Phylogeography of the Indo-Pacific parrotfish Scarus psittacus: Isolation generates distinctive peripheral populations in two oceans. Marine Biology, 157(8), 1679–1691. doi:10.1007/s00227-010-1442-4

Wong, B., & Candolin, U. (2015). Behavioral responses to changing environments. Behavioral Ecology, 26(3), 665–673.

Woodroffe, C. D., Brooke, B. P., Linklater, M., Kennedy, D. M., Jones, B. G., Buchanan, C., … Zhao, J. X. (2010). Response of coral reefs to climate change: Expansion and demise of the southernmost pacific coral reef. Geophysical Research Letters, 37(15), L15602. doi:10.1029/2010GL044067

Xue, A. T., & Hickerson, M. J. (2015). The aggregate site frequency spectrum for comparative population genomic inference. Molecular Ecology, 24(24), 6223–6240. doi:10.1111/mec.13447

