## Supplementary figures and images for "Coping with Pleistocene climatic fluctuations: demographic responses in remote endemic reef fishes"

### Figure S1

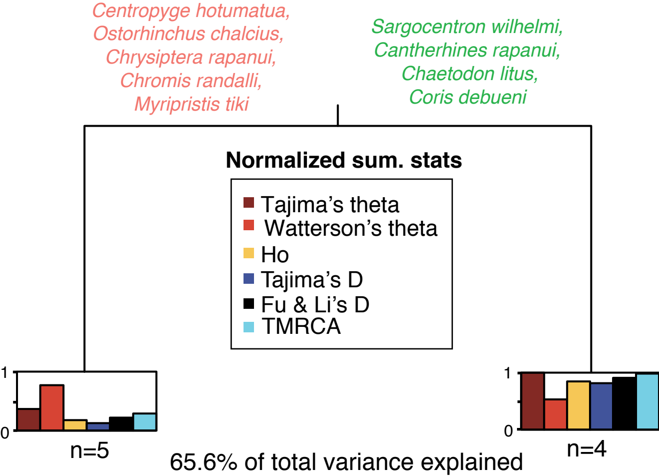

### Figure S2

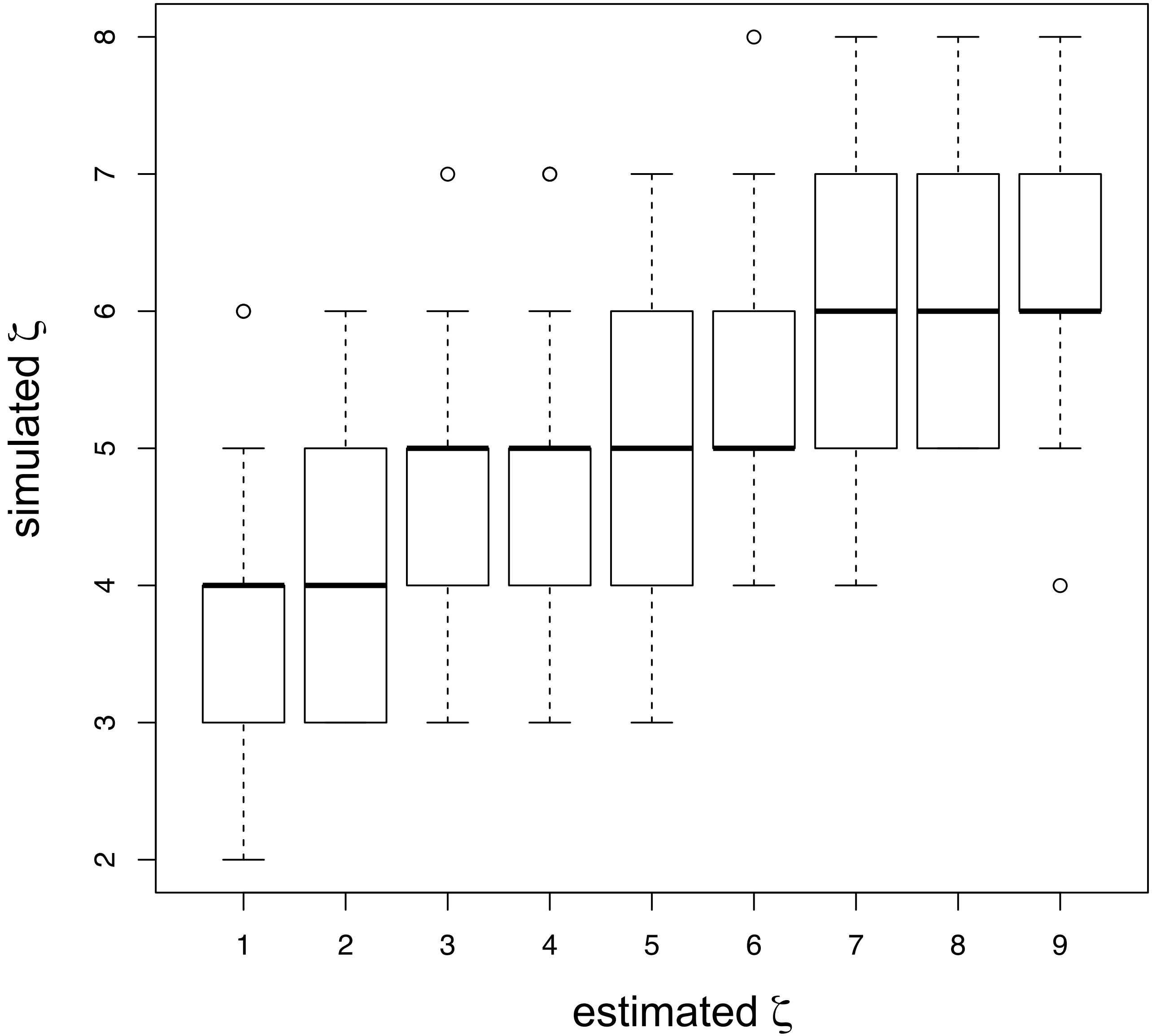

### Figure S3

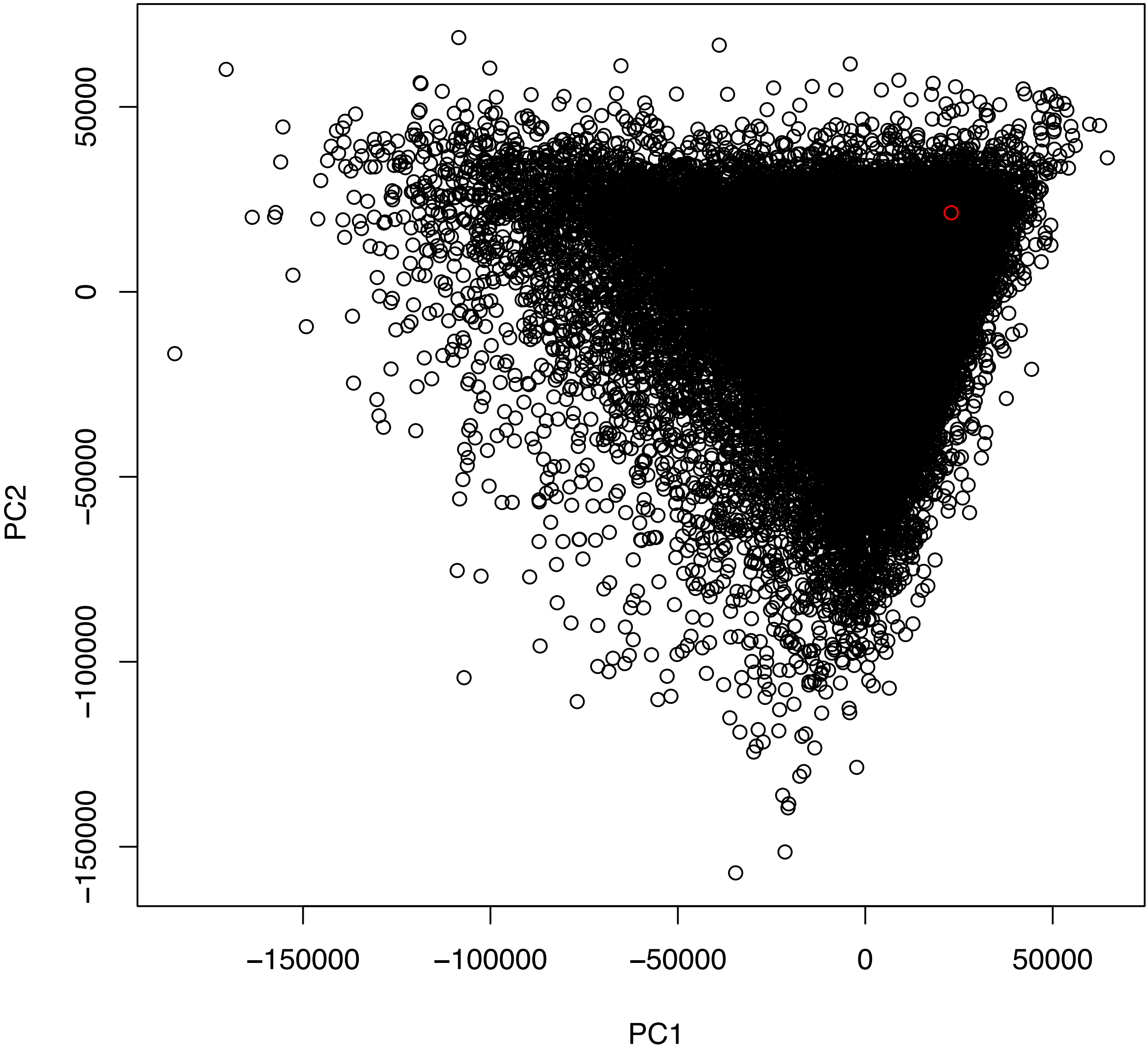
